# Spectral sparsification helps restore the spatial structure at single-cell resolution

**DOI:** 10.1101/2022.01.25.477389

**Authors:** Jingwan Wang, Shiying Li, Lingxi Chen, Shuai Cheng Li

**Author notes:** Corresponding authors Correspondence to Shuai Cheng Li. These authors contributed equally to this work (Jingwan Wang, Shiying Li).

## Abstract

Single-cell RNA sequencing thoroughly quantifies the individual cell transcriptomes but renounces the spatial structure. Conversely, recently emerged spatial transcriptomics technologies capture the cellular spatial structure but skimp cell or gene resolutions. Cell-cell affinity estimated by ligand-receptor interactions can partially reconstruct the quasi-structure of cells but falsely include the pseudo affinities between distant or indirectly interacting cells. Here, we develop a software package, STORM, to reconstruct the single-cell resolution quasi-structure from the spatial transcriptome with diminished pseudo affinities. STORM first curates the representative single-cell profiles for each spatial spot from a candidate library, then reduces the pseudo affinities in the intercellular affinity matrix by partial correlation, spectral graph sparsification, and spatial coordinates refinement. STORM embeds the estimated interactions into a low-dimensional space with the cross-entropy objective to restore the intercellular quasi-structures, which facilitates the discovery of dominant ligand-receptor pairs between neighboring cells at single-cell resolution. STORM reconstructed structures achieved shape Pearson correlations ranging from 0.91 to 0.97 on the mouse hippocampus and human organ tumor microenvironment datasets. Furthermore, STORM can solely *de novo* reconstruct the quasi-structures at single-cell resolution, *i.e*., reaching the cell-type proximity correlations 0.68 and 0.89 between reconstructed and immunohistochemistry-informed spatial structures on a human developing heart dataset and a tumor microenvironment dataset, respectively.

## Introduction

Revealing the spatial context and molecular abundance of cells and tissue is critical for understanding the composition and functions of complex tissues. Single-cell RNA sequencing (scRNA-seq) technologies quantify the single-cell transcriptome by a high sequencing depth with whole-transcriptome coverage^1^. The thorough scope of single-cell transcriptome enables investigations on cell heterogeneities, subpopulations, and interactions^2, 3^. However, the isolation procedure renounces the spatial context of these cells.

Spatial transcriptomics (ST) technologies have been developed to acquire spatial context and expression profiles simultaneously. High-plex RNA imaging technologies^4–6^ only localize dozens to hundreds of genes, and spatial barcoding technologies such as 10X Visium, Slide-Seq, and HDST^7–9^ yield a greater magnitude. However, they have achieved unsatisfied abundances or inadequate cell resolution, which restricts the potential of ST data for downstream analyses.

Except for wet-lab approaches, researchers also proposed computational methods to restore the spatial structure from the scRNA-seq data. NovoSpaRc^10^ assigns cells to tissue locations by probability. Its premise only considers the similarity in gene expression as the neighboring factor, neglecting the heterogeneity of, for instance, the transition areas^11^ or immune cell infiltration regions^12^. CSOmap reconstructs the intercellular proximity based on the contact-required ligand-receptor (LR) interactions^13, 14^. Specifically, CSOmap estimates the affinity of two cells by the mRNA expression summation of the interacting LR pairs, forming a *k*-nearest neighbor affinity graph simulating cell-cell interactions. However, the pseudo affinities in the affinity graph remain untended, leading to a defective reconstruction of spatial structure.

Researchers also started to integrate the ST data with the scRNA-seq data. Early attempts for integration focus on reconstructing cellular spatial structure based on spatial references such as immunohistochemistry (IHC) or fluorescence *in situ* hybridization (FISH)^15, 16^. Spatial barcoding presents a new aspect for integrating scRNA-seq and spatial data, leading to two primary integration approaches: deconvolution and mapping^17^. One objective of deconvolution methods is to infer the proportion of cell types from each ST capture location or spot in the ST data. Provided with a labeled scRNA-seq dataset, non-negative least squares and dampened weighted least squares linear regression can deconvolute the captured spot mixtures^18, 19^. Alternatively, deconvolution can be accomplished by fitting a model of negative binomial distribution or Poisson distribution to the scRNA-seq expression with the empirical data of ST spot as a prior. Subsequently, maximized posterior yields an estimation of the cell-type distribution^20–22^. Moreover, several studies on the tumor microenvironment (TME) map subgroups of single-cell to specific subregions in ST data by the enrichment score^23, 24^. These mappings improve the resolution on the subpopulation level but require prior clustering and annotation on both data types, which is inaccurate when mapping tissue regions comprised of mixed cell types. SpaOTsc^25^ maps cells by minimizing the gene expression dissimilarity between single-cell data and ST with the optimal transport distance, neglecting the heterogeneity in spot.

Here, we present a software package, STORM, that recapitulates the single-cell resolution cell quasi-structure of the spatial transcriptome from a sparsified affinity graph where the pseudo affinities are reduced by partial correlation^26^, spectral sparsification^27^, and spatial coordinates refinement. Instead of solely delivering cell-type acknowledgment, STORM locates single-cell expression profiles in spots from a candidate library, hence enabling the exploration of the spatial intercellular communication mechanisms at single-cell resolution.

## Results

### Overview of STORM algorithm: reconstructing spatial organization at single-cell resolution from the spatial transcriptome

STORM provides a prepossessing module for ST datasets which select and aggregate single-cell profiles representing the expression profile of each spot. For a spot of the spatial data, the module derives the quantities of each cell type by deconvoluted cell type proportions produced by the stereoscope^20^ and a prespecified parameter *ℓ*_*s*_ representing the average number of cells in a spot (Figure 1a). The module then aggregates a set of single cells agreeing with the derived quantities and maximizing the correlation between the aggregated cell expression profile and the ST spot. Note that if the paired single-cell data are unavailable, we can use a labeled single-cell candidate library of the similar tissue to create aggregations (Figure 1b).

**Figure 1.**
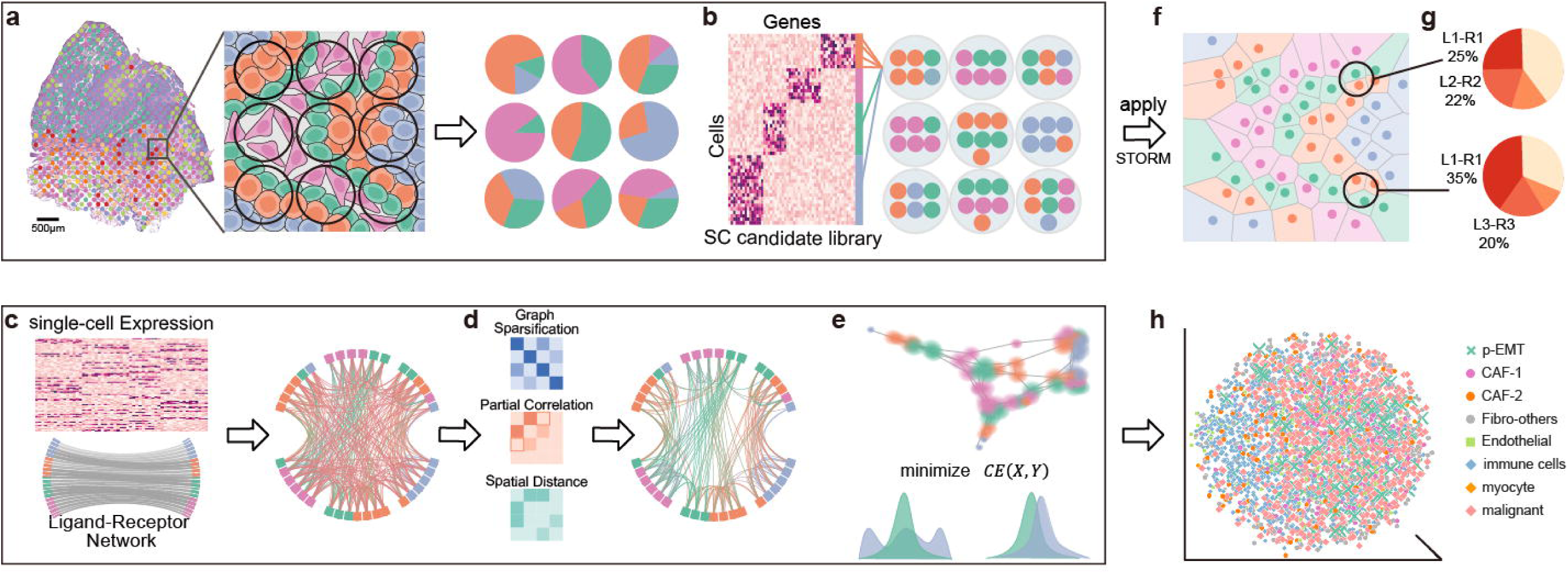
Schematics of STORM. **a-c**, Workflow of the preprocessing module. **a**, The preprocessing module of STORM adopts existing deconvolution software to decompose cell-type mixtures of ST profiles. **b**, The preprocessing module selects a designated amount of cells from the single-cell candidate library, equal to the estimated cell number per cell type in each spot. **c-e**, Workflow of the STORM. **c**, STORM derives the initial cell-cell affinity graph from the single-cell profiles by the LR interactions. **d**, STORM applies partial correlation, spectral graph sparsification, and spatial coordinates refinement on the cell-cell affinity graph to reduce pseudo affinities. **e**, STORM utilizes LFS embedding to embed interactions into a low-dimensional space. **f**, The 2D embedding of the selected single cells reconstructed by STORM. **g**, The determination of dominant ligand-receptor pairs between neighboring cells at single-cell resolution. **h**, The 3D embedding of the HNC data reconstructed by STORM.

Cells interact with proximal cells, and in this work, we use the term *affinity* as the measurement for the interaction strengths between interacting cells. We can build a cellular spatial configuration, termed *quasi-structure*, from the affinity values. We first assume that the cell-cell affinity can be estimated by the concentration of LR complexes which can be approximated by their mRNA abundance. Furthermore, we assume that cells compete for space because of the limitation of biological constraints. STORM has no prior knowledge of cell proximity when forming the initial affinity matrix. It calculates the affinity value between any two cells. Therefore, the approximated affinities based on the first assumption contain pseudo affinities between distant or indirect interacting cells. Following the above assumptions, STORM reconstructs the quasi-structure from scRNA-seq data with four steps: (a) establishing the initial affinity matrix by the LR expression profiles, which falsely includes the pseudo affinities between distant or indirect interacting cells; (b) constructing an affinity graph regards cells as vertices and the initial affinity matrix as the adjacency matrix; (c) reducing the underlying pseudo affinities in the initial affinity graph by partial correlation, spectral graph sparsification, and spatial coordinates refinement; and (d) embedding the sparsified affinity graph into a low-dimensional space as the quasi-structure in the single-cell resolution.

STORM approximates the cell-cell affinity by the mRNA abundance of interacting LR pairs (Figure 1c). For initial affinities of high variances, STORM replaces the initial cell-cell affinity matrix with the precision matrix to reduce the indirect correlations for subsequent procedures (Figure 1d). STORM reduces the pseudo affinities from the initial affinity matrix by imposing spectral graph sparsification and spatial coordinates refinement on the affinity matrix (Figure 1d). STORM adopts a local fuzzy set (LFS) embedding method to embed the processed affinity matrix to a low-dimensional space. The LFS step first builds a fuzzy topological representation from the processed affinity matrix, limiting the number of neighbors required by the second assumption (Figure 1e, top panel). Subsequently, the LFS step optimizes the representation in the low-dimensional space by minimizing the fuzzy set cross-entropy between the two representations (Figure 1e, bottom panel). STORM can take the curated single-cell aggregates, yielding the reconstructed quasi-structure for downstream analyses (Figure 1f). The embedding result, that is, the reconstructed quasi-structure by STORM, facilitates further evaluation of discovering dominant ligand-receptor pairs between neighboring cells at single-cell resolution (Figure 1g). Furthermore, with proper sparsification, STORM is capable of *de novo* reconstruction from the single-cell transcriptome. In the head and neck cancer (HNC) scRNA-seq dataset, STORM recapitulates the quasi-structure features which are commonly observed in the partial epithelial to mesenchymal transition (p-EMT) process: (a) p-EMT cells locating at the interface between malignant cells and cancer-associated fibroblasts (CAF) cells; (b) CAF-1 cells presenting at closer proximity to the p-EMT cells compared to CAF-2 cells; (c) malignant cells showing minimum interactions with immune cells due to immune evasion (Figure 1h).

### Assessing the performance of STORM in processing ST datasets

We demonstrate the performance of STORM on simulated and real-world ST datasets.

#### In silico *evaluation of STORM on processing the ST datasets*

We assess the validity of STORM in processing ST data coupled datasets by simulated datasets generated from the scRNA-seq data of the mouse hippocampus^28^. Since neurons, oligodendrocytes, and astrocytes are the main constituents of the hippocampus, we prepare two distinct scRNA-seq candidate libraries and coupled ST data for the simulated datasets: library A consisting of astrocytes and neuron cluster 1, and library B consisting of oligodendrocytes and neuron cluster 2. Moreover, the average number of cells per spot varies according to the tissue density and the spot diameter^23, 29, 30^. Therefore, we simulate ST data with the parameter number of cells per spot set as 10, 20, 30, and 40 to test the adaptability of the preprocessing module. Meanwhile, we perform five simulations for each parameter and candidate library to assess the robustness of STORM. Every simulated ST dataset consists of 30 spots. For each spot in the dataset, we arbitrarily sample the designated number of cells from each candidate library and regard the aggregated expression profile of these selected cells as the spot expression simulating the ST profile.

The preprocessing module of STORM selects 300 (*ℓ*_*s*_ = 10), 600 (*ℓ*_*s*_ = 20), 900 (*ℓ*_*s*_ = 30) and 1200 (*ℓ*_*s*_ = 40) cells respectively from each candidate library, constituting 30 single-cell aggregates to represent the expression profile of ST spots. The aggregated expression profiles of each single-cell aggregate regarding various parameters *ℓ*_*s*_ achieve an average Pearson correlation coefficient *r* = 0.97 with their corresponding ST profiles (Figure 2a). Moreover, we perform a paired t-test on the expression correlation of simulations across different cell-number parameters in each candidate library. In the best simulation of each candidate library, that is, the simulation with the highest average expression correlation, we observe that in candidate library A, the expression correlation differences between parameter ten and other parameters are significant. Yet, in candidate library B, the differences between parameters are not statistically significant (Figure 2b).

**Figure 2.**
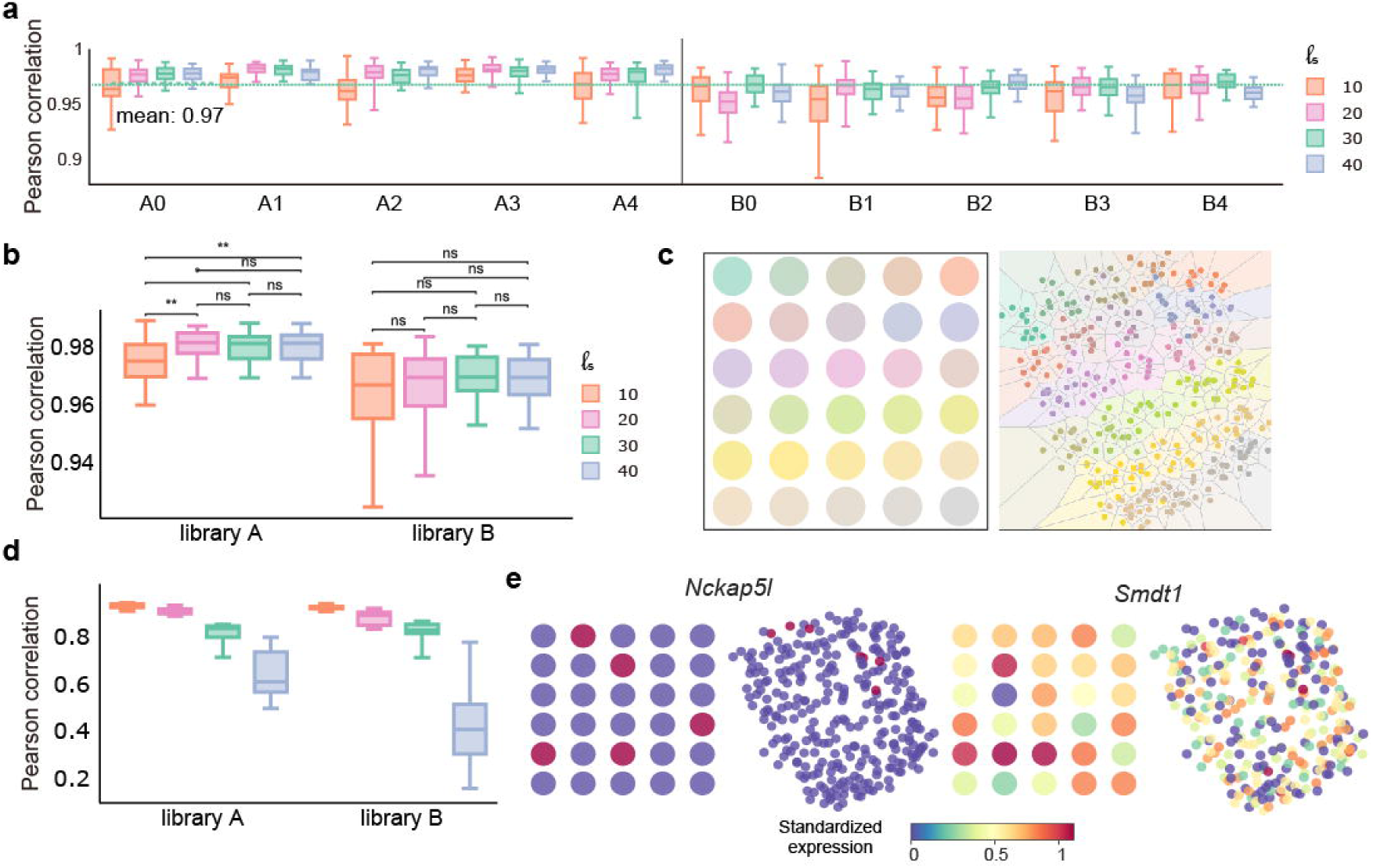
Evaluation of the validity and robustness of STORM on simulated datasets. **a**, The expression correlation between each spot and its corresponding single-cell aggregate regarding four cell-number parameters across five repeats of candidate libraries A and B. **b**, The best simulation of each cell-number parameter from libraries A and B, annotated with the statistical significance. Asterisks indicate level of statistical significance: ** - significance 0.01, * - significance 0.05, ns - not significant. **c**, The simulated ST structure (left) and the quasi-structure reconstructed by STORM (right). The color stands for each spot. **d**, The Pearson correlation of the pairwise distance between the reconstructed quasi-structure of STORM and its coupling ST spots. **e**, The standardized gene expression of exemplary genes in ST (left) and reconstructed quasi-structure (right).

Subsequently, STORM reconstructs the quasi-structure from the selected single-cell aggregates. The quasi-structure of each simulation reached a high shape correlation of *r* = 0.94, *ℓ*_*s*_ = 10 with the simulated spot organization (Figure 2c). The quasi-structure has a coincidental interspot organization as the cells originating from the same spot remain in the same compartment as illustrated by the Voronoi partition (Figure 2c, right). Furthermore, we compare the shape correlation between various parameters, and the average correlations decrease with the increase in cell number per spot (Figure 2d).

We also calculate the expression correlation of each gene between ST and single-cell aggregates. *Nckap5l* and *Smdt1*, achieve high expression correlations, *r* = 0.82, *r* = 0.67, between ST spots and single-cell aggregates (Figure 2e). The high-quality single-cell aggregates and the quasi-structure demonstrate the accuracy and robustness of the preprocessing module of STORM.

#### STORM reconstructed a high-quality quasi-structure for the mouse hippocampus dataset

We apply STORM to reconstruct the single-cell resolution quasi-structure for the mouse hippocampus dataset. The spatial data provided by stereoscope^20^ contains 609 spots, and the single-cell library from the *mousebrain.org* contains 8,449 cells, re-clustered and annotated by stereoscope, covering 56 subtypes across seven major groups.

The preprocessing module of STORM selects 6,071 cells that constitute 609 single-cell aggregates (*ℓ*_*s*_ = 10) from the single-cell candidate library to represent the expression profile of ST spots. Then STORM reconstructs the quasi-structure from the selected single-cell aggregates (Figure 3a). The reconstructed quasi-structure of STORM achieves a 0.97 Pearson correlation with its coupling ST spots in the pairwise distance (Supplementary information, Table S1).

**Figure 3.**
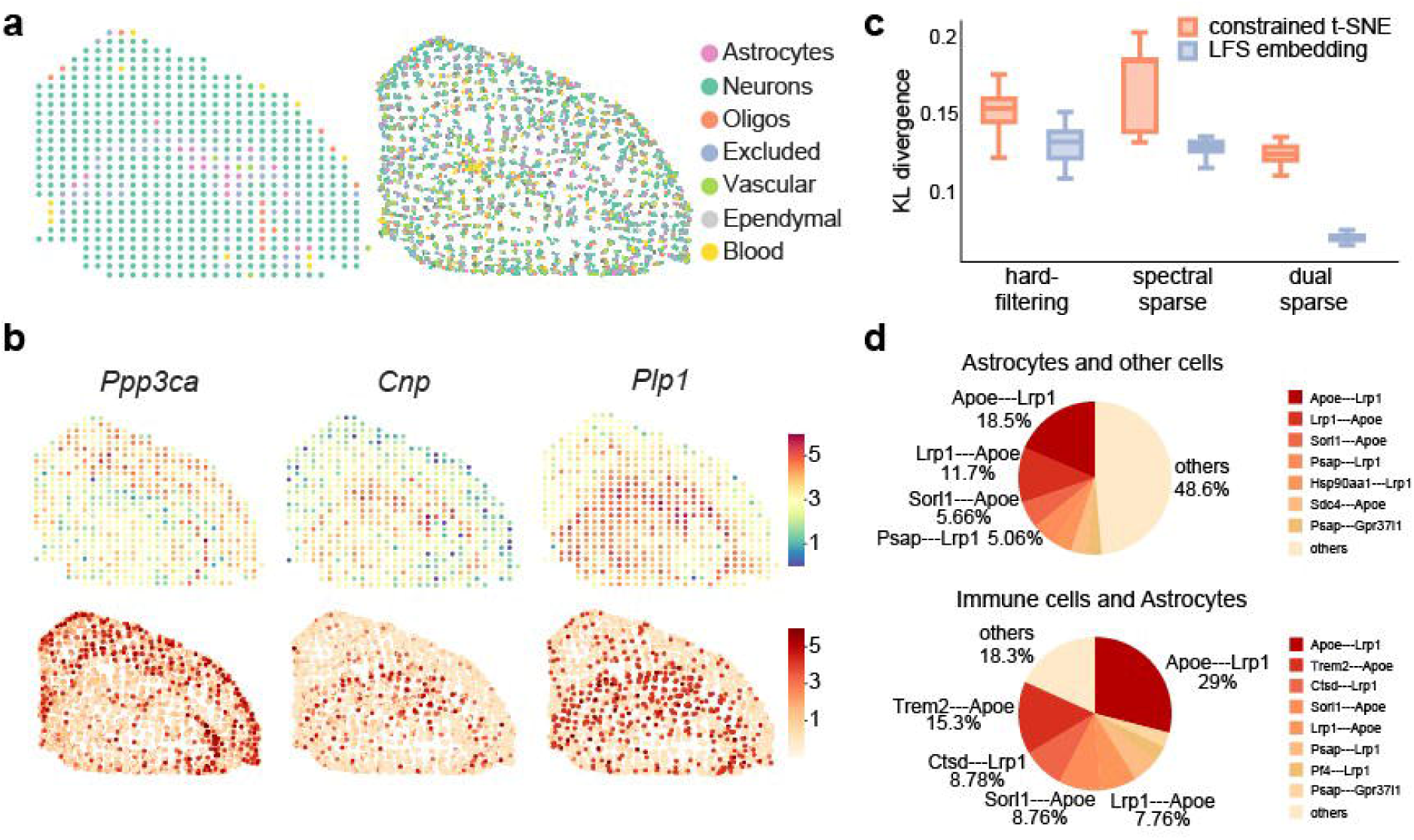
The reconstructed quasi-structure of mouse hippocampus. **a**, The 2D visualization of the ST spots (left) and the reconstructed quasi-structure of the mouse hippocampus (right), colored by cell types. **b**, The standardized gene expression of exemplary genes in ST (top) and reconstructed quasi-structure (bottom). **c**, The cell-type proximity KL divergence for the combinations of two different embedding methods and three sparsification methods. **d**, The pie charts of the LR pair contributions to the interactions of astrocytes with all other cells (top) and with only immune cells (bottom).

Furthermore, we calculate the expression correlation of each gene between ST and single-cell aggregates. *Cnp, Plp1* and *Ppp3ca*, achieve high expression correlations, *r* = 0.71, *r* = 0.70, *r* = 0.65, between ST spots and single-cell aggregates (Figure 3b, Supplementary information, Table S3). Meanwhile, the aggregated expression profile of each single-cell aggregate achieves a median Pearson correlation coefficient *r* = 0.66 with their corresponding ST profiles (Supplementary information, Fig. S1).

The cell-type proximity summarized by cell locations is vital for downstream analyses. Thus, the recapitulation of such information should also be a metric for evaluating the reconstructed quasi-structure. Specifically, we use Kullback-Liebler (KL) divergence to assess the difference of the cell-type proximity between the original and quasi-structure. The quasi-structure achieves a low KL divergence, 0.067, in the cell-type proximity (Figure 3c, Supplementary information, Table S2). Moreover, we assess the effectiveness of each step in STORM by comparing the KL divergence with different combinations of embedding and sparsification methods (Figure 3c). Comparing the LFS embedding that STORM utilizes with constrained t-SNE used by CSOmap, the lower median KL divergence in the combination of LFS embedding with a sparsification method is demonstrated. For sparsification methods, spectral graph sparsification partially reduces the pseudo affinities in the cell-cell affinity matrix, hence achieving a smaller median KL divergence compared to the hard-filtering method of keeping the top fifty high-affinity edges for each node. The additional distance metric provided by spatial information effectively reduces more pseudo affinities in the cell-cell affinity graph, leading to a smaller median KL divergence. The smallest KL divergence, 0.067, is acquired in the combination of LFS embedding and dual sparsification, which suggests the validity of each step in STORM.

The well-captured neighboring information in the reconstructed quasi-structure enables identifying the driver LR pairs mediating interactions between cell types. In the reconstructed quasi-structure, we observe that the interactions between lipoprotein receptor-related protein 1 (*Lrp1*) and apolipoprotein E (*apoE*) is the leading interactions among neurons, vascular cells, and astrocytes (Figure 3d, Supplementary information, Fig. S2). LRP1 mediates the metabolism of Amyloid-beta (*Aβ*), whose accumulation is a vital pathogenic element of Alzheimer’s disease. Yet apoE can block the LRP1-mediated pathway in astrocytes, hindering the clearance of *Aβ*^31^. Hence, certain immunotherapy targeting apoE has been applied on APP/PS1 mice to meliorate the accumulation of *Aβ*^32^. The reveal of the fundamental interaction between Lrp1 and apoE in our quasi-structure consolidates the validity of STORM and, therefore, its capability of providing valuable biological insights.

#### STORM uncovers the metastasis-promoting effect of HMGB1-SDC1 interaction in the human squamous cell carcinoma dataset

High-quality reconstructions of STORM help reveal the underlying molecular mechanisms of human diseases. We apply STORM on the human squamous cell carcinoma (SCC) dataset of patient 02 in Andrew *et al*.’s work^24^. The ST and scRNA-seq data are collected from the same malignant skin tissue. The spatial data contains 666 spots, and the matching scRNA-seq data contains 2,689 cells across 14 cell types.

The preprocessing module of STORM curates 6,625 cells with replacement regarding the SCC scRNA-seq data as the candidate library (*ℓ*_*s*_ = 10), forming single-cell aggregates to represent the expression profile of 666 spots in the spatial data. Subsequently, STORM rebuilds the quasi-structure from the curated single-cell aggregates. The reconstructed quasi-structure has high consistency, *r* = 0.91, with its coupling ST spot structure, regarding the pairwise distance (Figure 4a, Supplementary information, Table S1).

**Figure 4.**
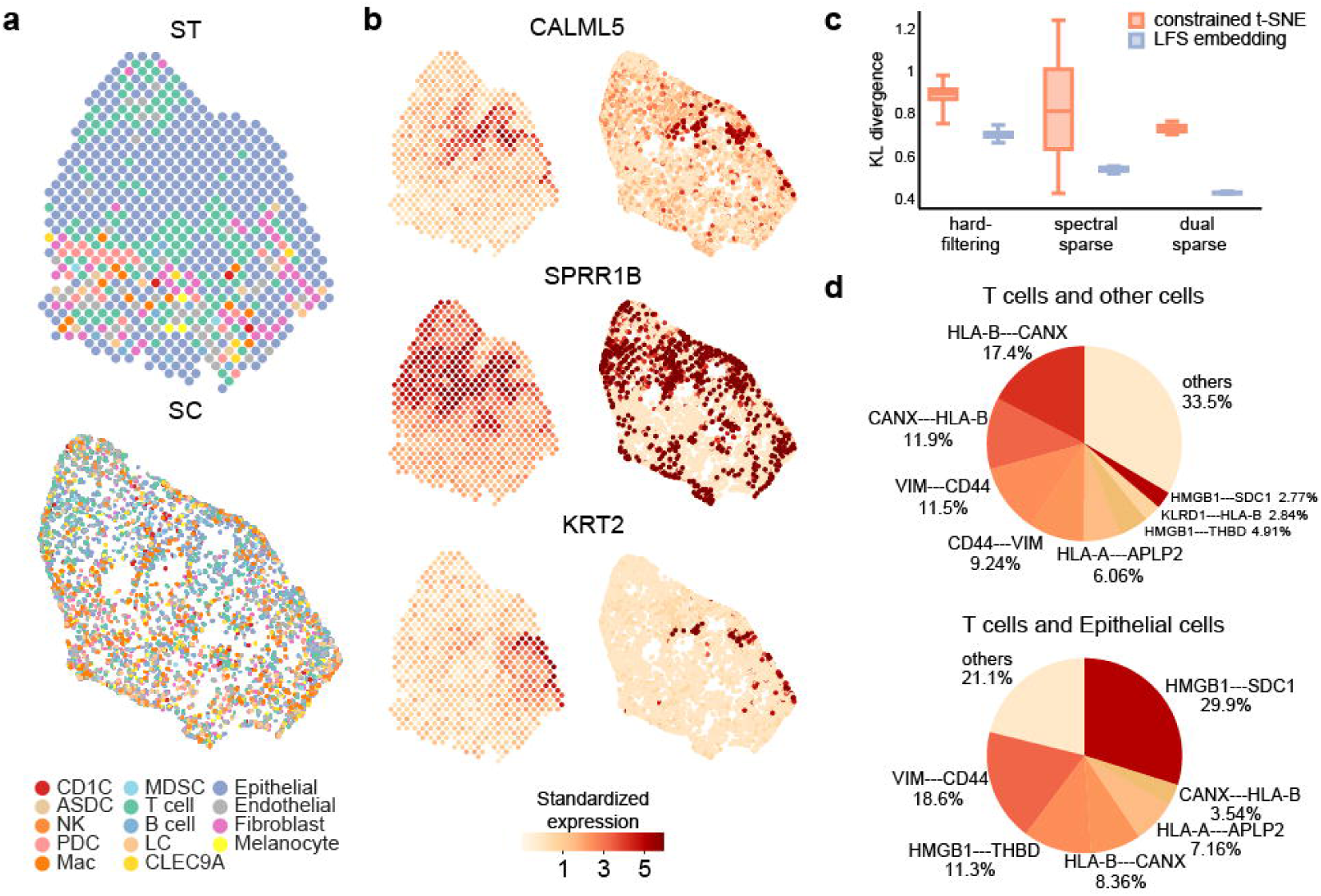
Performance of STORM on recapitulating the quasi-structure of SCC dataset. **a**, Spatial (top) and reconstructed quasi-structure (bottom) visualization of SCC, labeled by cell type. **b**, The standardized expression of cell-type marker genes in ST (left) and reconstructed quasi-structure (right). **c**, The cell-type proximity KL divergence of combining two embedding methods and three sparsification methods between ST and reconstructed quasi-structure. **d**, The pie charts of the LR pair contributions to the interaction of T cell and other cells (top), T cell and epithelial cells in particular (bottom).

Furthermore, we calculated the expression correlation of each gene between ST and single-cell aggregates (Supplementary information, Table S3). Several cell-type marker genes annotated in Andrew *et al*.’s work, *e.g*., *CALML5, SPRR1B, KRT2*, achieve high expression correlations, *r* = 0.79, *r* = 0.65, *r* = 0.61, between ST spots and single-cell aggregates (Figure 4b). Moreover, the aggregated expression profile of each single-cell aggregate has a median Pearson correlation coefficient *r* = 0.72 with their corresponding ST profiles (Supplementary information, Fig. S1).

The quasi-structure achieves a low KL divergence of 0.42 in the cell-type proximity between the original and the quasi-structure (Figure 4c, Supplementary information, Table S2). When comparing across different combinations of embedding and sparsification methods, STORM also achieves the smallest median KL divergence while combining dual sparsification and LFS embedding, which emphasizes the stability of STORM on cancer datasets.

Leveraging the high-quality quasi-structure STORM reconstructed, we identify the LR pair HLA-B-CANX as a driving force behind the interaction of T cells, constituting about 29% of the T cell affinities. Our finding is supported by a report regarding an impaired CD8+ T cell-mediated immune response due to the disturbance in HLA-B-CANX interaction in colorectal cancer^33^. We investigate the dominating LR pairs facilitating the crosstalk between T and epithelial cells. We identify that the interaction between HMGB1 and SDC1 contributes around 30% to the affinity between T and epithelial cells. HMGB1 and SDC1 have been reported to associate with the drug resistance in glioma^34^. Furthermore, the increase in HMGB1 promotes tissue invasion and metastasis of cancer^35^, and SDC1 influences the migration of mouse keratinocytes^36^. Our finding connects HMGB1 with SDC1, indicating that the reported promotion of metastasis may result from the interaction between HMGB1 and SDC1. The discovery demonstrates that the high-quality quasi-structure reconstructed by STORM facilitates disclosing the decisive LR interaction underneath the cell-cell communications.

#### STORM reveals different dominating LR pairs in two types of cancer cells from the high-quality quasi-structure

Tumor heterogeneity has been an obstacle to cancer therapy since mutant clones escape and thrive from the targeted therapy. Our spatially informed single-cell transcriptome can characterize the driver interactions between distinct subpopulations. We apply STORM on the patient PDAC-A of the pancreatic ductal adenocarcinoma (PDAC) dataset in Moncada *et al*.’s work^23^. Three tissue sections of PDAC-A were sequenced. We use the spatial data of replica 1. The ST and scRNA-seq data are processed from the same malignant tissue. The spatial data contains 428 spots, and the scRNA-seq data contains 1,926 cells annotated by 17 cell types.

Regarding the PDAC scRNA-seq data as the candidate library, the preprocessing module of STORM curates 4,289 cells with replacement (*ℓ*_*s*_ = 10), constructing single-cell aggregates to represent the expression profile of 428 spots in the spatial data. Given the high variance in the affinity values of the PDAC dataset, STORM reconstructs the quasi-structure of the curated single-cell aggregates with the precision matrix form of affinity matrix. The reconstructed quasi-structure achieves high similarity, *r* = 0.93, of the pairwise distance with its coupling spatial data (Figure 5a, Supplementary information, Table S1).

**Figure 5.**
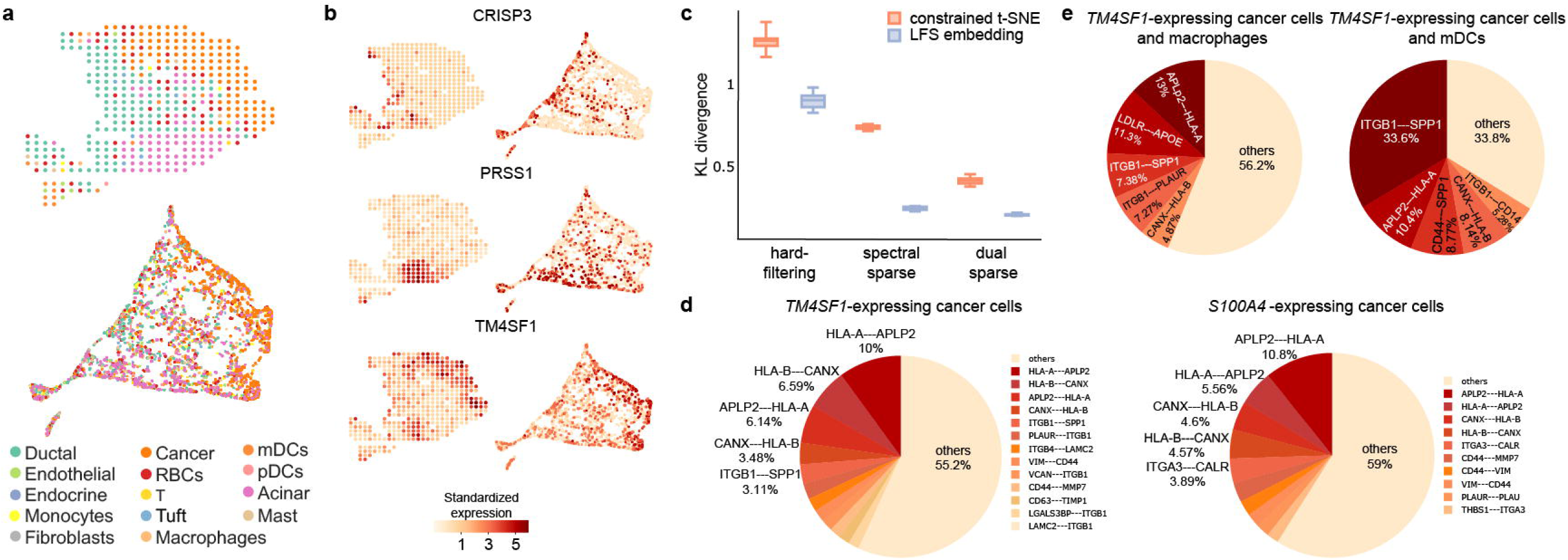
Performance of STORM in rebuilding quasi-structure from PDAC dataset. **a**, ST (top) and reconstructed quasi-structure colored by all cell types (bottom). **b**, The standardized expression of three genes in ST (left) and single-cell aggregates (right). **c**, The cell-type proximity KL divergence for the combination of three sparsification methods and two different embedding methods. **d**, The pie chart of LR pair contributions in *TM4SF1* – and *S100A4* -expressing cells. **e**, The pie chart of LR pair contributions between *TM4SF1* -expressing cells with macrophages (left) and mDC (right).

Moreover, we calculated the expression correlation of each gene between ST and single-cell aggregates (Supplementary information, Table S3). The feature gene of the main regions identified in Moncada *et al*.’s work, *CRISP3, PRSS1, TM4SF1*, also express in the corresponding regions in the quasi-structure (Figure 5b).

Furthermore, the quasi-structure achieves a low KL divergence of 0.13 in the cell-type proximity between the original and the quasi-structure (Figure 5c, Supplementary information, Table S2). The KL-divergence between ST and reconstructed quasi-structure decreases after progressively reducing the pseudo affinities by spectral graph sparsification and spatial coordinates refinement. The smallest median KL divergence is also achieved with the combination of LFS embedding and dual sparsification.

Subsequently, by evaluating the LR contribution to the cell-cell affinity, we observe that the interaction between HLA-A and APLP2 contributes around 16% to the overall interaction potential in both *TM4SF1*- and *S100A4* -expressing cancer cells (Figure 5d). APLP2 can cause a reduction in the expression of the total cell surface major histocompatibility complex (MHC) class I^37^, which is a crucial molecule for cancer cell recognition and elimination. The high interaction between HLA-A and APLP2 observed in the quasi-structure indicates a potential immune escape mechanism adopted by both *TM4SF1* - and *S100A4* -expressing cancer cells. Expect for the mutual LR interactions, we also found distinct dominating LR pairs in these two cancer types (Figure 5d). The LR pair ITGB1-SPP1 is a major contributing factor to the interaction between *TM4SF1* -expressing cancer cells between myeloid dendritic cell (mDC) and macrophage (Figure 5e). SPP1 has been proved to abet immune escape in lung adenocarcinoma through its mediation on macrophage polarization^38^. Experiments have also revealed how ITGB1-SPP1 interaction incites the cancer progression in ovarian cancer^39^. Our finding suggests that the interaction between ITGB1 and SPP1 potentially triggers the immune escape of PDAC. However, in *S100A4* -expressing cancer cells, the interaction between ITGA3 and CALR is more prevalent (Figure 5d, right). ITGA3 has been identified as a biomarker for diagnosing and prognostic predicting pancreatic cancer^40^. The LR pair ITGA3-CALR has also been predicted as a poor-prognostic LR pair by other datasets from the same tissue in the recent work of Suzuki *et al*.^41^. These discoveries demonstrate that researchers can characterize the tumor heterogeneity with the high-quality quasi-structure by revealing the driver interactions between distinct subpopulations.

### Evaluating the effectiveness of STORM on *de novo* reconstruction of single-cell datasets

We have demonstrated that the quasi-structure can be reconstructed from cell-cell affinity with proper sparsification. Therefore, we further evaluate the validity of STORM in reconstructing the spatial organization of scRNA-seq data without prior spatial structure.

#### STORM outperforms CSOmap on the hepatocellular carcinoma (HCC) dataset

We apply STORM on the HCC dataset consisting of 1,329 cells from Ren *et al*.’s work, for which the reconstruction of CSOmap obtains a Spearman correlation of *r* = 0.69 in the cell-type proximity with the IHC image of the same tumor sample. Given the large variance in the initial affinity values of the HCC dataset, STORM rebuilds the quasi-structure with the precision matrix form of affinity matrix. Compared with CSOmap, the reconstructed quasi-structure of STORM is visually less compact (Figure 6a) and achieves higher cell-type proximity, that is, a Spearman correlation of *r* = 0.89, with its IHC image reference (Figure 6b).

**Figure 6.**
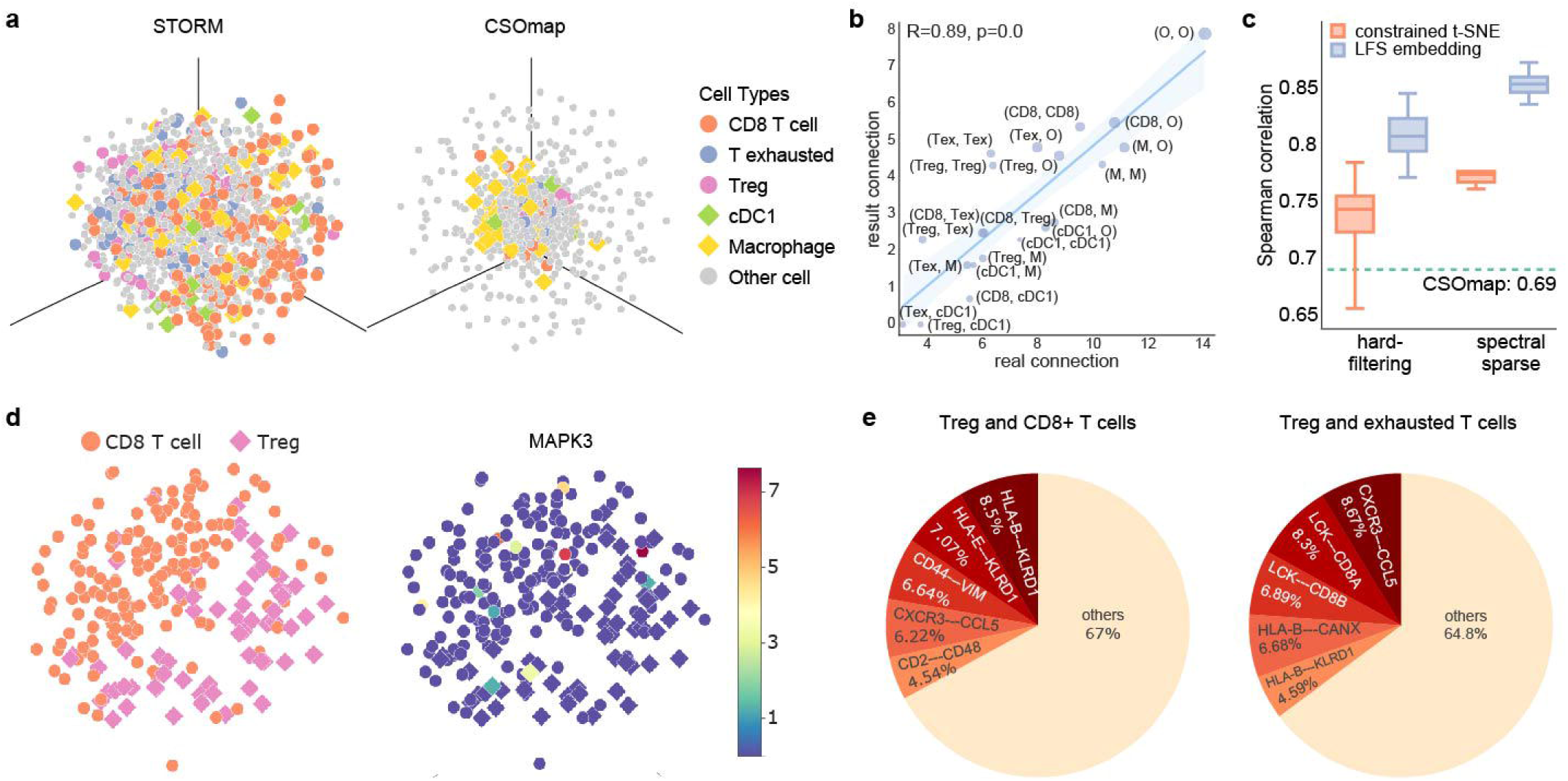
Application of STORM in restoring the quasi-structure of HCC. **a**, The 3D embedding of the reconstructed quasi-structure of STORM (left) and the prediction of CSOmap (right) on the HCC scRNA-seq data. **b**, Spearman correlation between IHC image-based cell connections (X-axis) and STORM reconstruction (Y-axis). CD8: CD8+ T cells; Tex: exhausted T cell; Treg: Foxp3+ regulatory T cells; M: macrophages; cDC1: CLEC9A+ dendritic cells; O: other cells. **c**, Comparison of cell-type proximity (Spearman) between different embedding and sparsification methods. The green dotted line represents the best Spearman correlation of the CSOmap prediction. **d**, CD8+ T cell and Treg cells in the quasi-structure of STORM colored by cell types (left) and standardized expressions (right). **e**, The pie charts of dominating LR pairs in the interaction of regulatory T cells with CD8+ T cells and exhausted T cells, respectively.

Subsequently, we evaluate the performance of combinations in embedding and sparsification methods regarding the cell-type proximity similarity (Figure 6b, Supplementary information, Table S4). A higher correlation is observed in the combination of LFS embedding and any sparsification method when comparing LFS embedding with constrained t-SNE. Moreover, spectral graph sparsification reduces the pseudo affinities, achieving a higher correlation than the hard-filtering method. The comparison between different combinations reveals the collaborative contribution of LFS embedding and spectral graph sparsification for reconstruction.

The high-quality reconstructed structure enables investigations on intercellular regulatory mechanisms. The interaction between regulatory T cells (Tregs) and CD8+ T cells suggests an ongoing suppression of the immune response^42^, during which Treg cells induce the p38 and ERK1/2 signaling pathways in effective T cells, which initiate DNA damage, resulting in cell senescence^43^. Consistent with the previous study, we observe an increase in the mRNA expression of ERK1 in the Treg-CD8+ T cell interacting area, indicating the potential of STORM in discovering the immune response signals hidden in the scRNA-seq data.

Furthermore, the well-captured cell-type proximity in the quasi-structure enables the analysis of the dominating LR pairs contributing to the cell-cell affinity. We analyze the main LR pairs between any two cell types. Specifically, we identified the difference in the dominating LR pair between Tregs and CD8+ T cells as well as between Treg and exhausted T cells, which indicates a distinct regulation mechanism of Treg in these two types of cells.*CCL5* is one of the signature genes identified in exhausted T cells^44^. The contribution of CXCR3-CCL5 increases in the interaction between Treg and exhausted T cells compared with CD8+ T cells. Indicating that the Tregs originated expression of CXCR3 may trigger the exhaustion.

The discovery demonstrates that the high-quality quasi-structure reconstructed *de novo* by STORM promotes the reveal of the LR interaction underneath the cell-cell regulatory mechanism.

#### STORM recapitulates the signal transmission process in the developing human heart

We apply STORM on a human developing heart dataset consisting of 3,717 cells from the 6.5 post-conception weeks (PCW) heart^45^. We apply STORM to reconstruct the quasi-structure of the heart dataset. The 3D quasi-structure of the developing human heart demonstrates a compact structure (Figure 7a, left). The atrial cardiomyocytes are spatially segregated from ventricular cardiomyocytes (Figure 7a, middle), which is consistent with the separation of the atrium and the ventricle in anatomy (Figure 7a, right). Moreover, we evaluate the cell-type proximity similarity between the quasi-structure and the *in situ* sequencing data. The quasi-structure achieves a high normalized Spearman correlation of *r* = 0.68 in the cell-type proximity.

**Figure 7.**
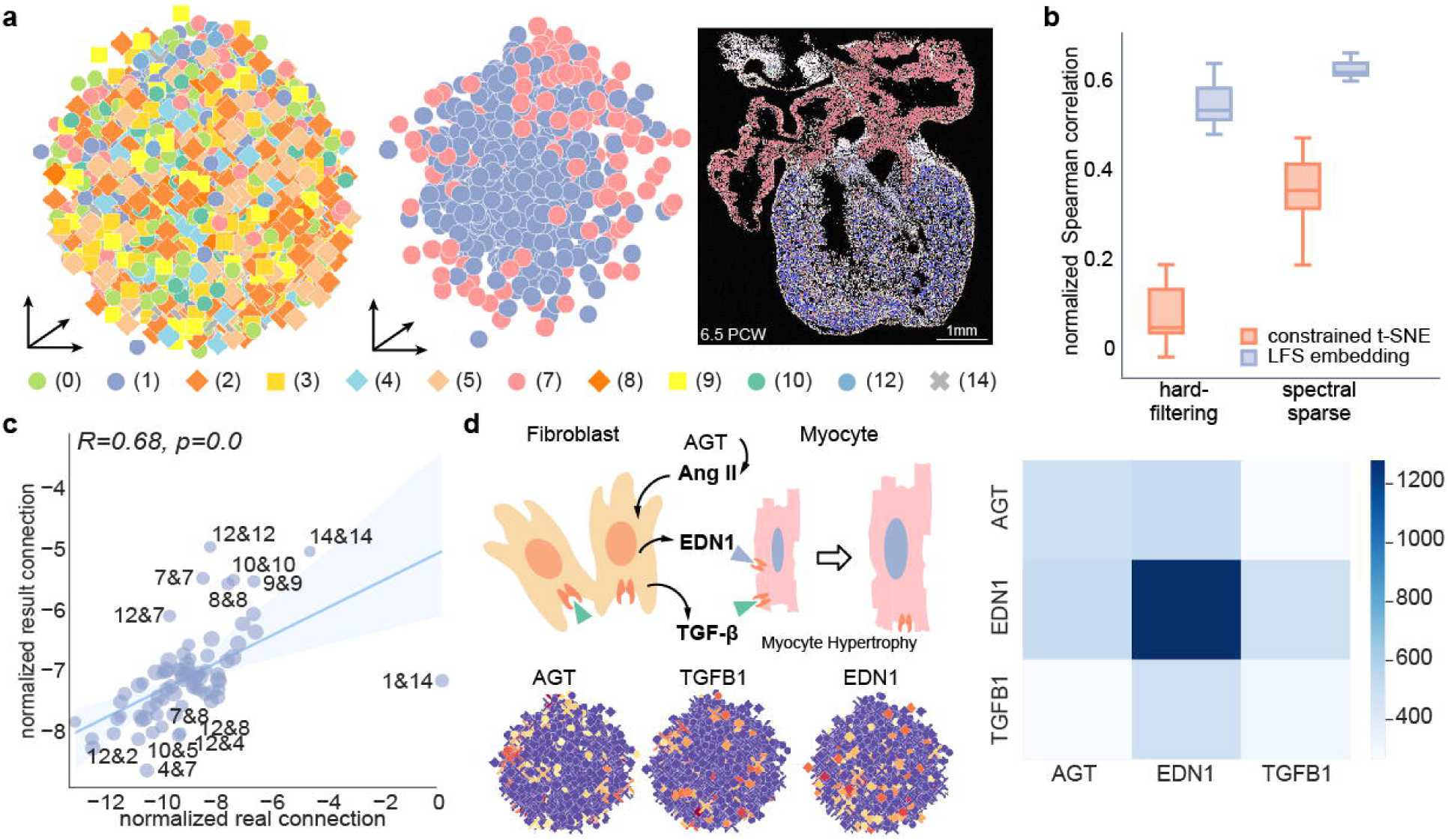
STORM recapitulates the quasi-structure of the developing human heart. **a**, 3D visualization of the reconstructed quasi-structure of developing human heart (left). Ventricular and atrial cardiomyocytes are separately displayed (middle). The tissue section of 6.5 PCW (scale bar: 1 mm), where the ventricular and atrial cardiomyocytes are manually labeled (right). Cell type label is the same as the original data: (0): Capillary endothelium; (1): Ventricular cardiomyocytes; (2): Fibroblast-like (related to cardiac skeleton connective tissue); (3): Epicardium-derived cells; (4): Fibroblast-like (smaller vascular development); (5): Smooth muscle cells / fibroblast-like; (7): Atrial cardiomyocytes; (8): Fibroblast-like (larger vascular development); (9): Epicardial cells; (10): Endohelium / pericytes / adventia; (12): Myoz2-enriched Cardiomyocytes; (14): Cardiac neural crest cells & Schwann progenitor cells. **b**, The normalized Spearman correlation of cell-type proximity in the result of hard-filtering and sparsified graphs embedded by constrained t-SNE (orange) and LFS embedding (blue). **c**, The normalized Spearman correlation between cell-type connections based on spots in the ST section (X-axis) and the quasi-structure reconstructed by STORM (Y-axis), with biases introduced by uneven cell counts among different cell types reduced after normalization. **d**, Mechanism illustration and evaluation of the regulation network between fibroblast and cardiomyocyte. Top-left: schematic diagram of molecular mediation between fibroblast and cardiomyocyte. Bottom-left: standardized expression of above intermediate genes. Right: heatmap of the numbers of neighboring pairs of cells expressing different marker genes.

When comparing the different combinations of embedding and sparsification methods, Figure 7b demonstrates that the reconstructed quasi-structure rebuilt by the combination of spectral sparsification and LFS embedding achieves the highest resemblance in cell-type proximity (Supplementary information, Table S4). The cell-type proximity STORM recapitulated includes fibroblasts and cardiac cells (Figure 7c), enabling fibroblasts to modify gene and protein expression, and ultimately cardiac function^46^. Ang II activates the paracrine secretion of TGF-*β*1 (*TGFB1*, transforming growth factor-*β*) and endothelin-1 (*EDN1*) in fibroblasts, leading to the cardiac myocyte hypertrophy (Figure 7d)^47^. Angiotensinogen (*AGT*) is a precursor for angiotensin I, which will be eventually converted to Ang II for further activities^48^. Therefore, we inspect the proximity of *AGT* high-express cell and *TGFB1, EDN1* high-express cell through the neighboring cell pair numbers between these cells in the quasi-structure(Figure 7d). We consider a pair of cells are neighboring if the distance is less than the median distance between any cell to its third-nearest neighbor. The proximity between cells that express critical signaling genes provides conditions for signaling through paracrine, consistent with the experimentally validated signaling pathway. This consistency indicates the effectiveness of the quasi-structure rebuild by STORM to reveal the local signal transmission process in the tissue.

## Discussion

The combination of the spatial context and expression profile of each cell enables our understanding of the intercellular regulation mechanism of tissue homeostasis and pathogenesis. The scRNA-seq discards the spatial context, and ST technologies skimp the cell resolution. Therefore, current technologies are inadequate to produce the spatial structure of tissues with single-cell resolution. In this work, we presented STORM to reconstruct the single-cell resolution spatial structure from the spatial and/or single-cell transcriptome. STORM rebuilds the quasi-structure of cells by embedding the sparsified affinity graph to a low-dimensional space. The reconstruction accuracy of STORM has been demonstrated in the mouse hippocampus, human heart, and tumor microenvironment of different organs in expression similarity, shape similarity, and cell-type proximity.

Although STORM relies on a comprehensive and valid LR pair database, extensive tests across different organisms and diseases demonstrate a consistent performance of STORM. The recapitulation of literature-supported major LR interactions in TMEs and immune responses also shows the effectiveness of the default LR datasets in providing valid biological observations. However, STORM can delineate a broader range of interactions with a higher accuracy if a more extensive LR pair network is expected with future developments. In addition, the prepossessing module benefits from a comprehensive single-cell candidate library. It is therefore subjected to the influence of sequencing depth of ST data, the imbalanced sizes, inconsistent cell-type constitution, and batch effects between ST and scRNA-seq data, and the accuracy of the estimated cell numbers per spot. Nevertheless, our evaluations consistently show that STORM produces high-correlation quasi-structures across various paired and unpaired datasets with different library sizes. In particular, we recommend using paired datasets for disease studies to ensure an accurate reconstruction against high heterogeneity among samples. In contrast, unpaired datasets have little influence on normal tissues with smaller divergence in mRNA expression across different samples.

Previous deconvolution methods^18–22^ failed to achieve a single-cell resolution, integrative methods either fall short in dealing with heterogeneous tissue^10, 23, 24^ or omit single-cell datasets without spatial reference^25^, and LR-based reconstruction^13^ neglected the pseudo affinities of distant or indirect interacting cells. Unlike previous methods, STORM utilizes the single-cell transcriptome, spatial transcriptome, and LR interactions to reconstruct a quasi-structure of cells in single-cell resolution by a curated affinity graph. A limitation of the preprocessing module is that the actual number of cells in each spot varies according to spots and tissues. For instance, tissue like the lung, which contains many alveoli, leaves plenty of cavities in the tissue section^49^. Therefore hard to estimate the cell number in each spot accurately. For future development, we intend to include an algorithm for accurate quantification of cell numbers per spot by the high-resolution histological image of the tissue section.

STORM reconstructs the spatial structure in single-cell resolution, utilizing the spatial context of each cell. The quasi-structure facilitates the acquisition of the dominating LR pairs in each cell pair, leading to the discovery of subpopulations based on dominating LR since cell talk subdivides cell functions. With a precise reconstruction, STORM reveals the co-occurrence of different types of cells and divergent colonization of subpopulations, which cannot be detected solely by scRNA-seq or ST technologies. Besides, STORM can acquire the dominating LR pairs in each cell pair, leading to the discovery of novel subpopulations based on dominating LR since cell talk subdivides cell functions. These abilities shed light on the studies on tumor heterogeneity and immune therapy. For instance, identifying the disparity of immune microenvironment around different cancer subpopulations could guide medication and metastatic evaluation. Furthermore, the quantification of intercellular interactions between the cancer cell and immune cell can predicate the prognosis of patients with clinical information.

## Materials and Methods

### Constructing the single-cell aggregates to reproduce ST expression profiles

We propose a preprocessing module to integrate ST data with scRNA-seq data. The module takes two parameters, the cell number *ℓ*_*s*_ and cell proportion *p*_*s,t*_, *t* ∈ *T* for a ST spot *s, T* denotes the set of cell types. The parameter *ℓ*_*s*_ denotes the average number of cells in a spot. The number of cells captured in a spot varies according to sequencing methods and tissue density; our module allows users to specify it. The software package *stereoscope*^20^ can infer *p*_*s,t*_. Stereoscope assumes a negative binomial distribution model on single-cell and ST data, building a reference expression profile of each cell type from the scRNA-seq data, then maximizing the posterior estimation to obtain the approximate cell proportion at every spot of the ST dataset.

Let *k*_*s,t*_ denote the cell number of type *t* at ST spot *s*, then *k*_*s,t*_ ≈ *ℓ*_*s*_ × *p*_*s,t*_ = *f*_*s,t*_. Note that *f*_*s,t*_ can be fractional. Here, we round on *f*_*s,t*_ randomly^50^ to acquire the integer number of *k*_*s,t*_ while stabilizing the expectation of *ℓ*_*s*_. Denoting the decimal part of *f*_*s,t*_ as {*f*_*s,t*_} ∈ [0, 1), *f*_*s,t*_ randomly rounds up or down to *k*_*s,t*_ according to the probability *P* (*k*_*s,t*_ = ⌈*f*_*s,t*_⌉) = {*f*_*s,t*_}.

The prepossessing module chooses cells from a predefined library to reproduce the single-cell resolution for the ST data. The summed expression profile of all chosen cells in **M**_**s**_ termed the *aggregated expressions E*(**M**_**s**_). It curates the single-cell aggregates set **M**_**s**_ by maximizing the Pearson correlation between *E*(**M**_**s**_) and the expression *E*(*s*) of spot *s*; that is, by the following objective function.

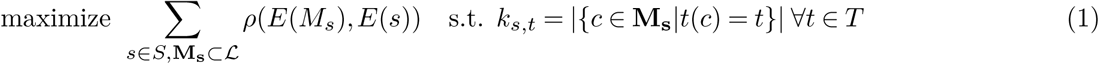

The number of chosen cells from each type in **M**_**s**_ should be the same as the value of *k*_*s,t*_. where *ℒ* ∈ ℝ ^*m*×*n*^ is the expression matrix of the single-cell library composed of *m* cells and *n* genes.

The module adopts a heuristic method of two steps, initialization and *swapping* to optimize the objective function. The initialization selects top *k*_*s,t*_ cells of type *t* for spot *s* according to the Pearson correlation coefficients between the spot and the cell from the single-cell candidate library.

If a better objective value is obtained, the swapping step swaps a cell in aggregates with a cell from the library. The process is repeated until convergence, or a predefined maximum number of iterations is achieved. The swapping process can be time-consuming, and we adopted a local sensitive hash (LSH) strategy to accelerate the swapping step^51^. During the swapping procedure, the module removes one cell from the aggregate **M**_**s**_ at spot *s* randomly, denoting the aggregate after the removal as **M**_**s**_′. The module chooses a new cell **m** in each iteration to further increases the *ρ*(*E*(**M**_**s**_′ ∪ {**m**}), *E*(*s*)). It can be chosen by querying a cell in LSH that has the highest correlation with *E*(*s*) − *E*(**M**_**s**_′).

The module performs feature selection^52^ on the single-cell candidates to reduce the noise introduced by sequencing and low variable genes by choosing the top 3,000 highly variable genes and 80% highly variable LR genes to maintain the capability to infer the intercellular affinity.

### Measuring the intercellular affinity by ligand-receptor interactions in single-cell profiles

We denote the single-cell expression matrix as **T** ∈ ℝ^*r*×*n*^ consisting of *r* cells and *n* genes. With *n*_*lr*_ ligand-receptor (LR) pairs, we define the ligand and receptor expression matrices as **T**_*L*_ and 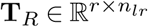, whose columns are the corresponding LR pairs’ ligand and receptor expressions, respectively. The multiplication of the two expression matrices yields the affinity between each pair of cells suggested by the co-expression of each LR pair. As a cell can simultaneously express both ligand and receptor genes, we have two symmetric terms 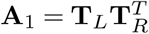 and 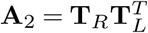 representing two possible LR orders in each cell pair. We formulate the *initial affinity matrix* **W** as 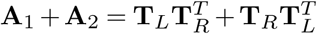 of size *r* × *r*.

### Reducing the pseudo affinities to refine the affinity matrix by sparsification

The initial affinity matrix includes pseudo affinities between distant or indirectly interacting cells. Here we present three different approaches for diminishing the pseudo affinities, that is, partial correlation, spectral graph sparsification, and spatial coordinates refinement for ST coupled datasets.

We first adopt partial correlation to reduce the pseudo affinities for initial affinities of high variance^26^. Partial correlation identifies the latent variables representing direct causation and removes indirect relationships among entities^53, 54^. While a covariance matrix represents the relations between any two entities, the inverse of a covariance matrix, also known as the precision matrix, approximates the partial correlations among entities^55^. For the block expression matrix 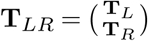, we denote its covariance matrix as **K**, that is, 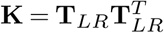. In particular, we have the block form of 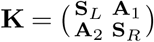, where 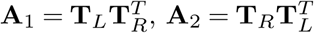, and 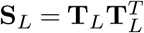 and 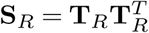 represent the ligand and receptor gene expression similarity between any two cells. We could distinguish direct and indirect LR interactions among cells and keep the direct ones by using the precision matrix of **K**, *i.e*., 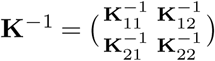. Therefore, we have 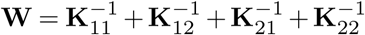 representing direct LR interactions.

We build the affinity graph **G** by regarding the cells as vertices and the cell-cell affinity as the edge weight. When the context is clear, we also refer **W** as the adjacency matrix for **G** for notation simplicity. We further denote the Laplacian matrix of **G** as **L**. Therefore, we apply the Spielman-Srivastava spectral graph sparsification algorithm^56^ to remove pseudo affinities. Spectral graph sparsification aims to find a sparse approximation of the original graph while maintaining high spectral similarity between two graphs^27^. In the Spielman-Srivastava algorithm, the effective resistance, *i.e*., the distance between two vertices connecting by an edge is proportional to the reciprocal of its edge weight. In the sparsification step, edges are sampled by the probabilities proportional to their effective resistances. The algorithm preserves the spectrum of the graph Laplacian, *i.e*. the eigenspaces spanned by eigenvalues, and their relations by requiring high similarity between the two Laplacian matrices, while some previous works only maintain the span of the dominant eigenvectors^57, 58^. We define the effective resistance between two cells *u* and *v* as

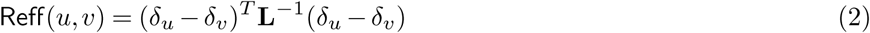

where *δ*_*u*_ ∈ {0, 1}^*r*^ is the indicator vector of vertex *u*. Following the definition, the sparse graph preserves the crucial edges of the original graph. We sample the edge (*u, v*) by the probability *p*_*u,v*_ = *min*{1, *C* · (log *r*)*W*_*u,v*_ · Reff(*u, v*)/*ϵ*^2^}, where *C* is some constant and *ϵ* is the approximation parameter. We further adjust the weight of the sampled edge (*u, v*) as *W*_*u,v*_/*p*_*u,v*_. We determine the value of the term *C*/*ϵ*^2^ by the user-defined proportion of preserved edges *q* = 2 ∑ *u,v p*_*u,v*_/*r*(*r* − 1). Since the expected number of chosen edges can be bounded by

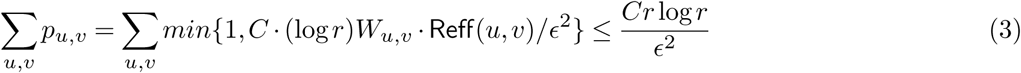

where 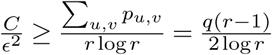, thus by adjusting the parameter *q* we can control the percentage of preserved edges. Moreover, we utilize the spot coordinates in the coupled spatial data as one sparsification approach. If two cells belong to nonadjacent spots, the affinity between them is considered to be pseudo affinities.

### Reconstructing the quasi-structure with fuzzy set cross-entropy embedding

The embedding of a cell-cell affinity graph to a low-dimensional space consists of two stages: (a) forming a topological representation 𝕎 of sparsified the cell-cell affinity **W**; and (b) finding an embedding 𝔼 in the low-dimensional space of the topological representation to minimize the discrepancy between the embedding and the representation. A reliable topological representation of **W** should maintain the affinity relations while restricting the number of neighbors for each cell. Here, we maintain the top *k*_*n*_ affinities in **W** for each cell while setting other values to be zeros. Subsequently, we perform min-max normalization on the remaining affinities to obtain the membership strength in the range of [0, 1], denoting the matrix as 𝕎. The fuzzy simplicial set expands the classical binary definition of membership by allowing continuous membership strength in the range of [0, 1]^59^, and the union of the fuzzy simplicial sets^60^ yields the fuzzy topological representation. Hence, 𝕎 is the fuzzy topological representation of **W**.

Subsequently, we apply strategies from UMAP^61^ to minimize the fuzzy set cross-entropy between the embedding 𝔼 and the topological representation 𝕎, that is,

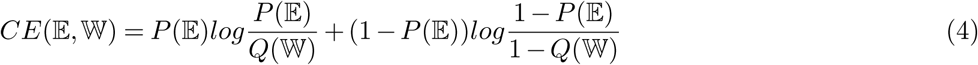

where *P* (𝔼) and *Q*(𝕎) represent the normalized adjacency matrices of 𝔼 and 𝕎, respectively. We use a spectral layout, that is, the Laplacian matrix of 𝕎 to as the initial Cartesian coordinates of 𝔼^62^. By regarding edges as attractive forces and vertices as repulsive forces, we alternatively apply the attractive and repulsive forces until *CE*(𝔼, 𝕎) converges to a local minimum.

### Evaluating the reconstruction performance of STORM

A major metric for assessing the quality of the reconstructed spatial structure is its reproduction of the spatial characteristics of the tissue. Given a spatial structure of cells, we construct a fixed-volume neighbor graph, where the radius is the median distance between any cell to its third-nearest neighbor. According to the fixed-volume neighbor graph, we quantify the spatial characteristics as the number of neighboring pairs between any two cell types, indicating whether the two are enriched or depleted near each other. Therefore, we evaluate the cell type enrichment or depletion discrepancy by the Kullback-Leibler (KL) divergence^63^ of the neighboring pair numbers for any two cell types between a given spatial structure and the embedding structure. To further evaluate the statistical significance of observed possible enrichment or depletion, we compare the number of neighboring pairs with 1000 random permutations of the cell type labels. We test the enrichment hypothesis, that is, the observed number of neighboring pairs is larger than the random expectation by *p*-values from both the right-tailed and left-tailed tests. We further adjust the *p*-values following the Benjamini-Hochberg procedure^64^ and obtain the *q*-values with a cutoff of 0.05 for significance.

### Revealing the dominating LR pairs contributing to intercellular affinity

Given a pair of cell expression profiles **E**_*i*_ and **E**_*j*_, the contribution from the *k*-th LR pair to the total cell-cell interacting affinity can be formulated as:

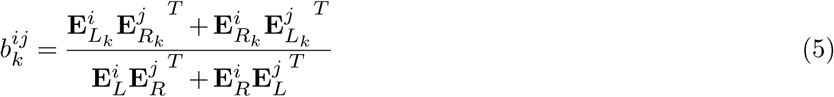

The contribution of each LR pair between two cell types *t*_1_ and *t*_2_ is calculated by:

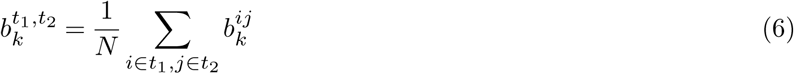

Where *N* is the number of neighboring cell pairs between *t*_1_ and *t*_2_.

## Data availability

The ST and scRNA-seq data we use has been previously published^13, 20, 23, 24, 28, 45^ and are available online mousebrain.org, https://github.com/almaan/stereoscope. The PDAC, HNC, and SCC datasets are deposited at the Gene Expression Omnibus under GSE111672, GSE103322, and GSE144240. The count matrix of the developing human heart is available at https://www.spatialresearch.org with the erythrocytes and immune cells removed, and the labels we use in this work remain consistent with the original publication. The HCC dataset CSOmap used is deposited at EGA with accession number EGAS00001003449.

## Code availability

The software implementation and analysis notebooks of STORM are available at https://github.com/deepomicslab/STORM.

## Acknowledgements

We would like to express sincere gratitude to Dr Wenji Ma for the suggestions on data collection. We appreciate Ms Wenqian Zhang for the manual cell-type annotation of the heart tissue section. This project was supported by SIRG (CityU SIRG 7020005).

## Author Contributions

These authors contributed equally: Jingwan Wang, Shiying Li

## Contributions

SCL conceived and designed the project. J.W. developed the software. J.W. and SYL performed the analysis and validation experiment. SCL, J.W., SYL, and L.C performed manuscript writing, review, and editing.

