## supplementary figs for "Spectral sparsification helps restore the spatial structure at single-cell resolution"

### Supplementary information, Fig. S1

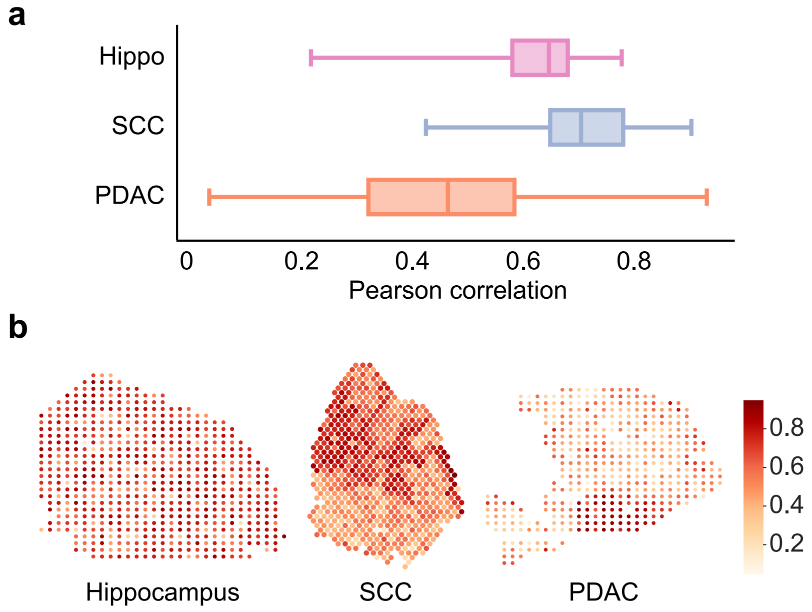


Fig. S 1: The expression correlation of each spot between spatial and single-cell aggregates in ST coupled datasets. **a**, The boxplot of gene correlation between each spot of ST and its corresponding single-cell aggregates in the hippocampus, SCC, and PDAC datasets. **b**, The heatmap of the expression correlation of each spot in the hippocampus, SCC, and PDAC datasets.

### Supplementary information, Fig. S2

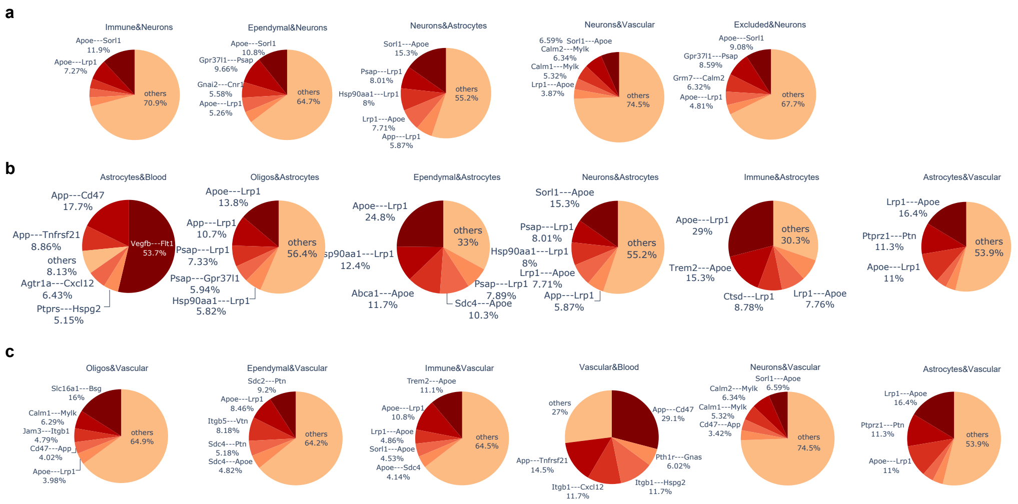


Fig. S 2: The LR contribution calculated from the reconstructed quasi-structure of the mouse hippocampus dataset. **a**, The pie chart of LR contribution between neurons and other cell types. **b**, The pie chart of LR contribution between vascular cells and other cell types. **c**, The pie chart of LR contribution between astrocytes and other cell types.
